# Dynamic Pathway of Guanidine-III Riboswitch Folding Revealed by Single-Molecule FRET: Mg^2+^-Assisted Preorganization and Ligand-Induced Kinetic Trapping

**DOI:** 10.1101/2025.04.15.648854

**Authors:** Mengke Jiang, Yunqiang Bian, Lujun Zou, Tingting Liu, Zongzhou Ji, Weitong Ren, Zilong Guo, Yi Cao, Yonghua Jiao, Hai Pan

## Abstract

Riboswitches are structured RNA elements that regulate gene expression by sensing and binding small molecules. The guanidine-III riboswitch, a critical bacterial regulator responding to guanidine toxicity, undergoes precise conformational changes that remain poorly characterized at a dynamic, mechanistic level. In this study, we employed single-molecule Förster Resonance Energy Transfer (smFRET) coupled with molecular dynamics (MD) simulations to delineate how the guanidine-III riboswitch transitions among distinct conformational states. We identify three principal states ― an extended (E-state), a compacted prefolded intermediate (P-state), and a folded pseudoknot structure (F-state) ― with rapid interconversion in the absence of ligand. Magnesium ions significantly stabilize intermediate states via a cooperative, preorganization mechanism, enhancing ligand binding efficiency. Binding of guanidine drastically suppresses the reverse transitions, kinetically trapping the riboswitch into its active folded state primarily through a conformational selection mechanism, with additional contributions from induced-fit dynamics. This work illuminates the unique dynamic pathway by which the guanidine-III riboswitch integrates ionic and ligand cues, ensuring precise gene regulatory responses in bacteria.

## INTRODUCTION

Riboswitches, non-coding structured RNA elements found within untranslated regions of mRNAs, regulate gene expression by selectively binding small metabolites(1-3). They serve as essential biological sensors, modulating gene expression in metabolic pathways, transport, and stress responses to maintain cellular homeostasis(4,5). Among these regulatory RNAs, the guanidine-III riboswitch, belonging to the ykkC family, stands out due to its role in sensing guanidine—an essential yet potentially toxic metabolite in bacteria(6,7).

Guanidine has dual roles: serving as a precursor in nitrogen-rich metabolites and as a cellular toxin at high concentrations by disrupting protein functions(8). Bacteria manage guanidine homeostasis by employing the guanidine-III riboswitch, typically positioned upstream of genes encoding transporters or enzymes that detoxify or export guanidine. Upon binding guanidine, the riboswitch adopts a folded pseudoknot structure, thereby altering downstream gene expression to counteract elevated guanidine levels(9).

Structural studies, particularly through crystallography, have revealed static views of riboswitch-ligand complexes but leave significant gaps concerning dynamic and intermediate conformations critical to their regulatory functions(10). Techniques such as smFRET have recently provided powerful insights by capturing transient conformations and real-time transitions, thereby addressing limitations inherent to ensemble-based methodologies(11,12).

In this investigation, we employed smFRET alongside MD simulations to provide unprecedented details of the conformational transitions and folding pathway of the guanidine-III riboswitch. We specifically elucidated how magnesium ions facilitate riboswitch folding by preorganizing intermediate states, thereby enhancing guanidine binding efficiency through cooperative interactions. Furthermore, we characterized how guanidine binding acts via a kinetic trapping mechanism—significantly suppressing reverse transitions and stabilizing the riboswitch in its biologically active folded conformation. Our findings thus clarify the dynamic pathways and mechanisms underlying riboswitch function, deepening the understanding of RNA-based gene regulation and highlighting potential targets for antimicrobial strategies.

## MATERIALS and METHODS

### RNA synthesis

The oligonucleotides of the guanidine-III riboswitch were synthesized by GenScript, Nanjing, China. The sequence was adopted from a previous structural study(13) with minor revisions: 5’-UCCGGACGAGGUGCGCCGUACCCGGUCAGGACAAGACGGCGCUUU-3’. In particular, an extra uracil was added at the 5’ end for Cy5 labeling, Cy3 was labeled in the middle at position U25 and with three extra Us appended at the 3’ end for biotin labeling.

### Chemicals and reagents

Magnesium chloride hexahydrate (MgCl_2_·6H_2_O), sodium chloride, glucose oxidase, catalase, and Trolox (6-hydroxy-2,5,7,8-tetramethylchroman-2-carboxylic acid) were purchased from Sigma-Aldrich. Tris-HCl (pH 8, 1 M stock solution), EDTA, guanidine, methylguanidine, aminoguanidine and urea were purchased from Sangon Biotech, Shanghai, China.

### Single-molecule FRET experiments

Single–molecule measurements were carried out on an Olympus IX–73 based objective–type total internal reflection fluorescence (TIRF) microscope equipped with an oil immersion UAPON 100xOTIRF objective lens (N.A. = 1.49, Olympus).

We assembled a microfluidic channel in between a clean glass slide and a PEGylated coverslip as described previously(14). The RNA sample was dissolved in 1X T50 buffer (10 mM Tris–HCl, pH 8, 50 mM NaCl), heated up to 85 °C for 3 mins, and then snap-cooled on ice. The RNA solution was diluted to an appropriate concentration and flowed into the microfluidic channel, and incubated for 3 min. The RNA molecules were immobilized on the passivated glass surface through biotin-streptavidin interaction. The unbound free RNA was washed away using imaging buffer (1xT50 with or without Mg^2+^/Gdm^+^). For the experiments without divalent metal ions, EDTA was added to the imaging buffer to a final concentration of 2 mM as a chelator. An oxygen scavenging system containing 0.8% (w/v) D-(+)-glucose, 1 mg/mL glucose oxidase, 0.04 mg/mL catalase, and 2 mM Trolox was introduced into the microfluidic channel to reduce fluorophore photobleaching and photoblinking(15). The emissions were split into two channels by a dichroic mirror (FF640–FDi01, Semrock) with two band–pass filters for the donor channel (ET585/65, Chroma Tech) and the acceptor channel (FF01–698/70, Semrock), respectively. A 532 nm laser (green) was used to excite Cy3, and emission from both Cy3 and Cy5 was collected by an EMCCD (iXon 897, Andor Technology) with a temporal resolution of 50 ms. To exclude the possibility that the vanishment of the acceptor signals was due to Cy5 photobleaching, a direct excitation of Cy5 using a 640 nm laser was briefly executed (∼2 s) at the end of the recording window. To obtain higher temporal resolution data, the bin size of the image was set to 2×2 and the ROI was set to 1/2, in which case the time resolution was increased to 10 ms by using a lower sample concentration.

### Single-molecule FRET data analysis

We processed raw movie files using IDL (Research Systems) to extract smFRET time traces that were analyzed using MATLAB scripts. The FRET efficiency was calculated using E_FRET_ = I_A_/(I_A_ + I_D_) and the distance between dyes was calculated using r = (1-E)^1/6^/E^1/6^·R_0_ with R_0_ value of 54 Å for Cy3-Cy5 pair(16). FRET efficiencies observed in each trace were combined to generate FRET distribution histograms using 10-frame average in Origin. For kinetic data, traces before photobleaching were manually selected for consistency, anticorrelated dye behavior, and single step photobleaching. Hidden Markov modelling software (HaMMy)(17) was then applied using a three-state model. Transition rates were generated using TDP software(17). TODPs(18), which depict the fraction of counts transition from a specific initial FRET state to a final FRET state among total FRET counts were generated using a Python program.

### Molecular dynamics simulations

An all-atom structure-based model was employed to investigate the conformational dynamics of the RNA riboswitch, with all heavy atoms explicitly represented. The reference structure for the native conformation was obtained from the Protein Data Bank (PDB code: 5nwq). The initial structures were generated using the SMOG2 package(19), and all simulations were conducted with the OpenMM MD engine(20), which integrates the OpenSmog library(21). Multiple unbiased simulations were conducted simultaneously using identical parameters but different seeds to sample a sufficient number of folding and unfolding events. Starting from an unfolded structure, simulations were executed using Langevin dynamics with an inverse friction constant of 1 ps. A time step of 0.002 time units (ps) was utilized, and each trajectory’s total simulation time amounted to 2 × 108 MD steps. The simulation temperature was set to 1.6 temperature unit, as frequent folding and unfolding events were observed.

## RESULTS and DISCUSSION

### smFRET Characterization reveals Three major conformational states of Guanidine-III Riboswitch in the absence of ligand

Recently, the structure of the ykkC guanidine III riboswitch from *Thermobifida fusca* was solved by X-ray crystallography(13). The 41 nt long riboswitch adopts an H-type pseudoknot structure formed by two interconnected stem-loops (P1 and P2), in which terminal loops base-pair reciprocally (Figure 1A). Its compact core, stabilized by extensive hydrogen bonding and non-Watson-Crick interactions, forms the guanidine-binding pocket between C6, G7 and G17 (Figure 1A, inset). To investigate the structural dynamics of the guanidine-III riboswitch, we employed single-molecule FRET (smFRET) under ionic conditions of 50 mM NaCl, pH 8. The riboswitch construct was immobilized on polyethylene glycol (PEG)-passivated, streptavidin-coated slides via a biotin labeled at the 3′-end of RNA (Figure 1B, C). To probe global folding transitions, we positioned fluorophores at sites optimized to report on pseudoknot formation: a Cy5 acceptor was appended to an extra uracil residue introduced at the 5′ end of the aptamer core, while a Cy3 donor was site-specifically conjugated to U25 within the non-conserved junction loop between P1 and P2 (Figure 1B). This labeling strategy ensures minimal perturbation of the pseudoknot interface while maximizing distance changes between fluorophores during folding.

**Figure 1.**
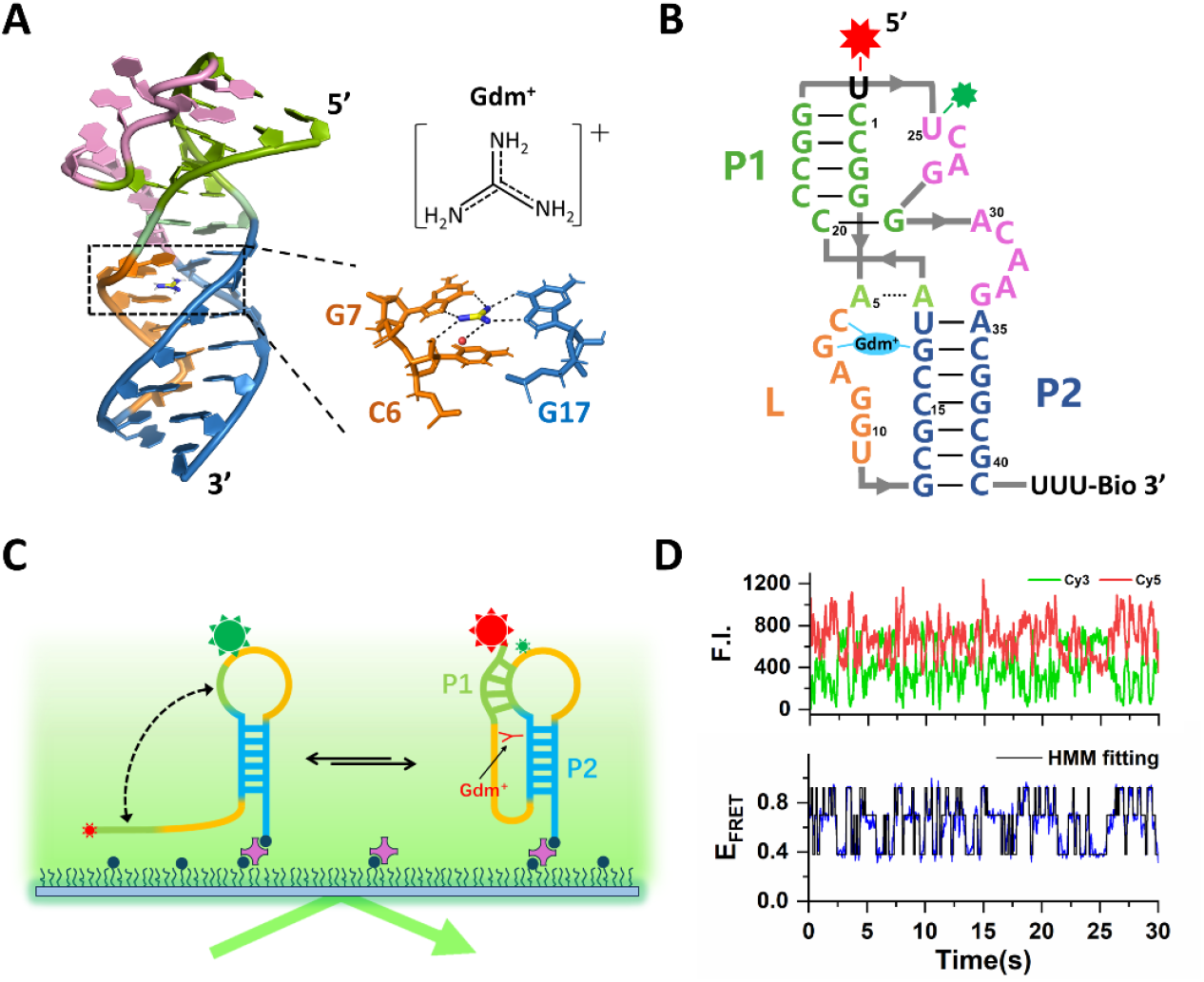
Scheme diagram of strategy for TIRF-based smFRET measurement of guanidine-III riboswitch. **(A)** Crystal structure of the *Thermobifida fusca* ykkC guanidine-III riboswitch (PDB: 5nwq). **(B)** Guanidine-III riboswitch sequence drawn in the pseudoknot secondary structure, comprising stem-loops P1 and P2, with fluorophore labeling present. **(C)** Schematic representation of the TIRF-based smFRET analysis of the folding of guanidine-III riboswitch. **(D)** Representative smFRET trace in 50 mM NaCl. Hidden Markov model (HMM) fitting showed in black lines.

As shown in Figure 1D, using hidden Markov modeling (HMM) to fit the FRET trajectories, we identified three discrete FRET states in the absence of Mg^2+^ and guanidine: a median-FRET state (E_FRET_ ≈ 0.7) and a high-FRET state (E_FRET_ ≈ 0.92), corresponding to a partially compacted intermediate (denoted “prefolded”, P-state), and a fully folded pseudoknot conformation observed in ligand-bound crystal structures (“folded”, F-state). The inter-dye distances for these two states were calculated to be ∼47 Å and ∼35 Å, respectively. These two conformational arrangements are consistent with other pseudoknot riboswitches explored by smFRET studies(22-25). Surprisingly, an even lower FRET state (E_FRET_ ≈ 0.38, ∼69 Å) was detected, reflecting an extended conformation (“extended”, E-state). These states were populated dynamically under equilibrium conditions, with frequent interconversions observed over time. The FRET population histogram showed the riboswitch primarily populated in the extended state (56.4%, Figure 2A) under conditions without Mg^2+^ or ligands. These observations suggest that the guanidine-III riboswitch exhibits intrinsic conformational plasticity, sampling prefolded and folded-like states independent of Mg^2+^ or ligand. To further validate the 3-state-transition, we generated the transition occupancy density plots (TODPs) in Supplementary Figure S3A, in which clearly showed that 3 subpopulations interconvert dynamically. It is worth noting that, a fully unfolded structure with E_FRET_ centered at ∼ 0.18 (Supplementary Figure S1) was observed in addition to the three conformational states described above. However, such traces only accounted for less than 3% (26 traces out of 1120). Therefore, we excluded data from unfolded state in subsequent analysis.

**Figure 2.**
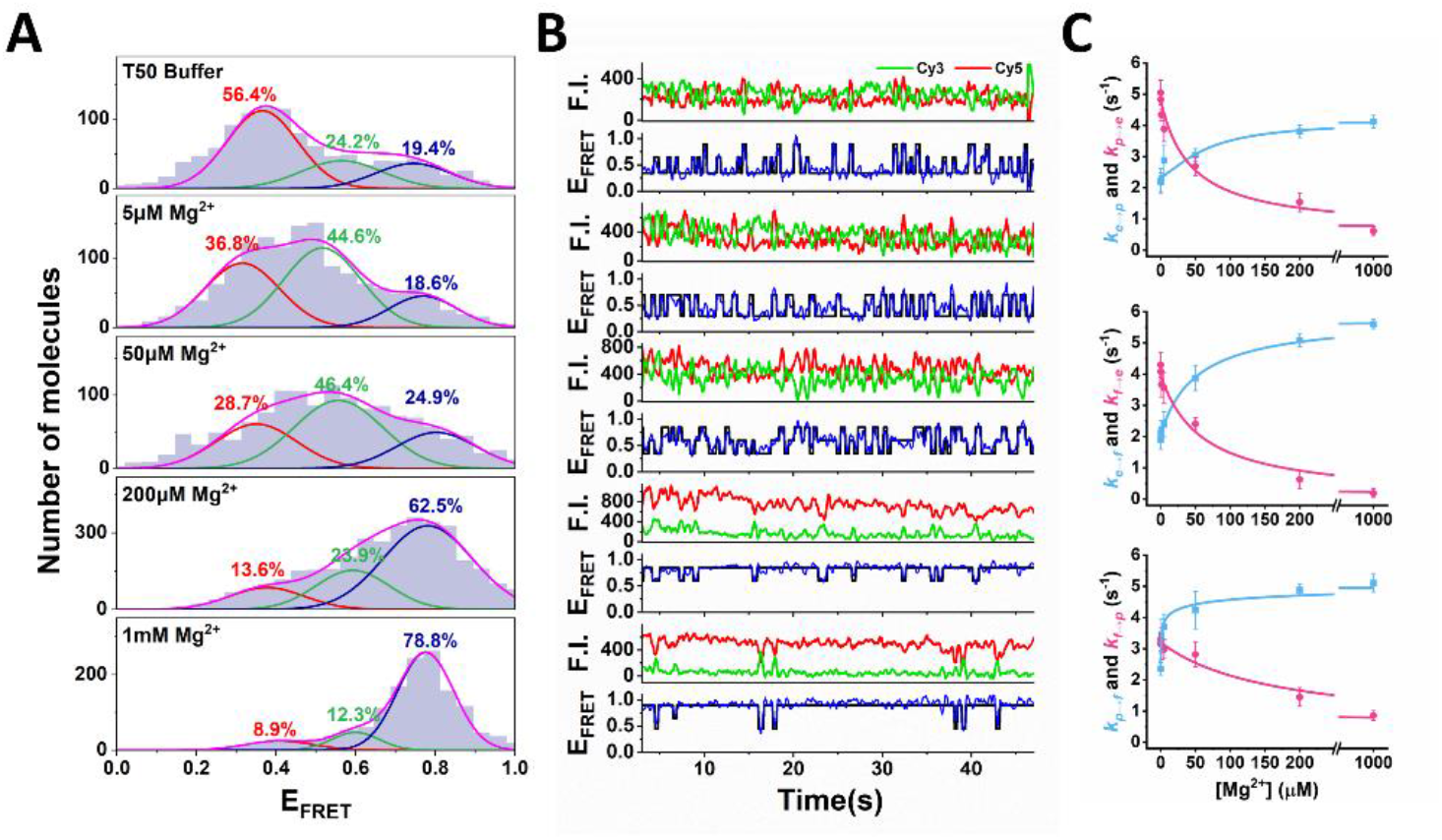
Mg^2+^ -dependent folding dynamics of the guanidine-III riboswitch by smFRET. **(A)** FRET population histograms at various Mg^2+^ concentrations. The fraction of three-state from Gaussian fitting is marked in the figure, red for extended state, green for prefolded state and blue for folded state, respectively. **(B)** Representative smFRET traces for guanidine-III riboswitch in different Mg^2+^ concentrations ranging from 0 to 1mM. The idealized FRET traces generated by the hidden Markov model (black) were overlaid on the experimental traces (blue). **(C)** Transition rate constants for guanidine-III Riboswitch in different Mg^2+^ concentrations.

In the crystal structure, the binding pocket forms a left-handed helical ramp through triple helix interactions with stem P2, even in the absence of guanidine, implying the structure of guanidine-III riboswitch might be highly dynamical. To test this hypothesis, we performed smFRET experiments using TE buffer without any salts (10mM Tris-HCl, pH 8, 2mM EDTA). As expected, the RNA aptamer exhibited fast extended-folded state transitions (Supplementary Figure S2A) resolved by higher temporal resolution recordings (10 ms, see MATERIALS and METHODS). The transition occupancy density plots confirmed two-state transition (Supplementary Figure S3B). We measured the transition rates by extracting the dwell time of each state fitted with hidden Markov model (HMM), which gave *k*_e→f_ = 8.3 s^-1^ and *k*_f→e_ = 11.2 s^-1^, respectively (Supplementary Figure S2B, C). The extended-folded state transition rates under the condition of 50mM Na^+^ were also extracted for a comparison. The values of *k*_e→f_ decreased to 1.9 s^-1^ and *k*_f→e_ decreased to 4.1 s^-1^ when sodium ions present. The reduced transition rates in the presence of monovalent ions (e.g., Na^+^) likely arise from electrostatic screening effects(26-29), which stabilize both the extended (E) and folded (F) states. Concurrently, these conditions prolonged the lifetime of the prefolded (P) intermediate state, enabling its experimental resolution through smFRET.

### Mg^2+^ stabilizes the prefolded state and facilitates the folded state of Guanidine-III riboswitch

Divalent magnesium ions (Mg^2+^) play a critical role in RNA structural integrity, serving as essential counterions that counteract the repulsive forces between phosphate groups and stabilize tertiary interactions vital for riboswitch function(30-33). To dissect how Mg^2+^ regulates the structural stability of the guanidine-III riboswitch, we performed smFRET experiments across a physiologically relevant Mg^2+^ gradient (from 100 nM to 5 mM) in the absence of guanidine. The average E_FRET_ values for E-state, P-state and F-state were ∼0.35, ∼0.6 and ∼0.85, respectively (Figure 2A, B).

Similar to the conditions in T50 buffer, the riboswitch was predominantly in the extended state at low Mg^2+^ concentrations (100 nM, E_FRET_ ≈ 0.35, 49.9%, Supplementary Figure S4A), intermittently sampling the intermediate/prefolded state (P-state) and folded state (F-state). As Mg^2+^ increased, the E-state occupancies declined concomitantly with a rise in the F-state population (Figure 2A). This transition aligns with previous observations of RNA folding, where Mg^2+^ acts as a bridging ion, stabilizing transient intermediates first and then promoting global folding(24,34,35). The Hill coefficient (n = 1.22) obtained from fitting F-state occupancy versus Mg^2+^ concentration further supported positive cooperativity, indicating that Mg^2+^ binding events are not independent but synergistically enhance RNA folding(36).

Notably, the P-state occupancy exhibited a biphasic response: an initial increase at submicromolar Mg^2+^ followed by a decrease at higher concentrations. This suggests that Mg^2+^ initially stabilizes the partially folded intermediate, which then acts as a precursor for folding. The F-state dominated (78.8%) at 1 mM Mg^2+^, indicating near-complete stabilization of the folded conformation.

To further elucidate how Mg^2+^ affects the intrinsic structural dynamics of guanidine-III riboswitch, we plotted the forward and reverse transition rates for each transition as a function of Mg^2+^ concentrations (Figure 2C). All three transitions showed similar trends, with forward rates (*k*_e→p_, *k*_e→f_, *k*_p→f_) increasing and reverse rates decreasing (*k*_p→ e_, *k*_f→e_, *k*_f→p_). Specifically, the forward rate increased by approximately 2∼3-fold (from *k*_e→p_p = 2.19 s^-1^ to 4.12 s^-1^ at 1 mM Mg^2+^ for E-P transition, from *k*_e→f_ =1.8 s^-1^ to 5.6 s^-1^ for E-F transition and from *k*_p→f_ = 2.36 s^-1^ to 5.11 s^-1^ for P-F transition), while the reverse rate dropped dramatically, from *k*_p→e_ = 5.05 s^-1^ to 0.61 s^-1^ (∼10-fold) for E-P transition, from *k*_f→ e_ = 4.3s^-1^ to 0.19 s^-1^ (∼20-fold) for E-F transition and from *k*_f→ p_ = 3.16 s^-1^ to 0.86 s^-1^ (∼4-fold) for P-F transition, respectively.

This bifurcation in mechanism is analogous to other riboswitches, where Mg^2+^ accelerates ligand recognition by preorganization of aptamer folding(37). Our data suggest that Mg^2+^ plays a similar role in the guanidine-III riboswitch, where partial folding intermediates are stabilized first to enable efficient folding. In particular, the increments in the forward rates towards folding and the reductions in reverse rates imply that the RNA folding facilitated by Mg^2+^ operates through both conformational selection and induced fit mechanisms(38,39).

### Effect of Gdm^+^ Ligand on Guanidine-III Riboswitch Structural Dynamics

Having established that Mg^2+^ ions stabilize the guanidine-III riboswitch in the folded conformation by modulating its conformational energy landscape, we next explored how the cognate ligand, guanidine (Gdm^+^), drives structural dynamics to enable metabolite-responsive gene regulation. Similar to Mg^2+^, Gdm^+^ shifts the riboswitch population towards a high-FRET state (F-state, E_FRET_ ≈ 0.8, Figure 3A, B), corresponding to the compact pseudoknot conformation observed in ligand-bound crystal structures(13). Titration of Gdm^+^ revealed a concentration-dependent increase in the folded fraction, fitting to a Hill equation with a cooperativity coefficient n = 0.92 and dissociation constant K_D_ = 57 μM (Supplementary Figure S4B), which is comparable to the in-line probing results(7). The near-unity Hill coefficient indicates non-cooperative binding, consistent with a single ligand interaction site, as previously validated in biochemical assays(7). Transition rate analysis showed similar decrements in reverse rates for all three transitions compared to Mg^2+^-only conditions (Figure 3C). The value of *k*_p→e_ decreased from 5.05 s^-1^ to 0.4 s^-1^ for extended-prefolded transition, *k*_f→e_ decreased from 4.3 s^-1^ to 0.14 s^-1^ for extended-folded transition and *k*_f→p_ decreased from 3.16 s^-1^ to 0.44 s^-1^ for prefolded-folded transition, respectively. However, the pronounced decelerations in reverse rates compared to Mg^2+^-only conditions suggest that Gdm^+^ binding suppresses interconversion between states, reflecting Gdm^+^-specific interactions within the ligand-binding pocket, which rigidify the pseudoknot core and disfavor partial unfolding. These reverse rates plateau at 500 μM Gdm^+^, indicating saturation of the kinetic trapping.

**Figure 3.**
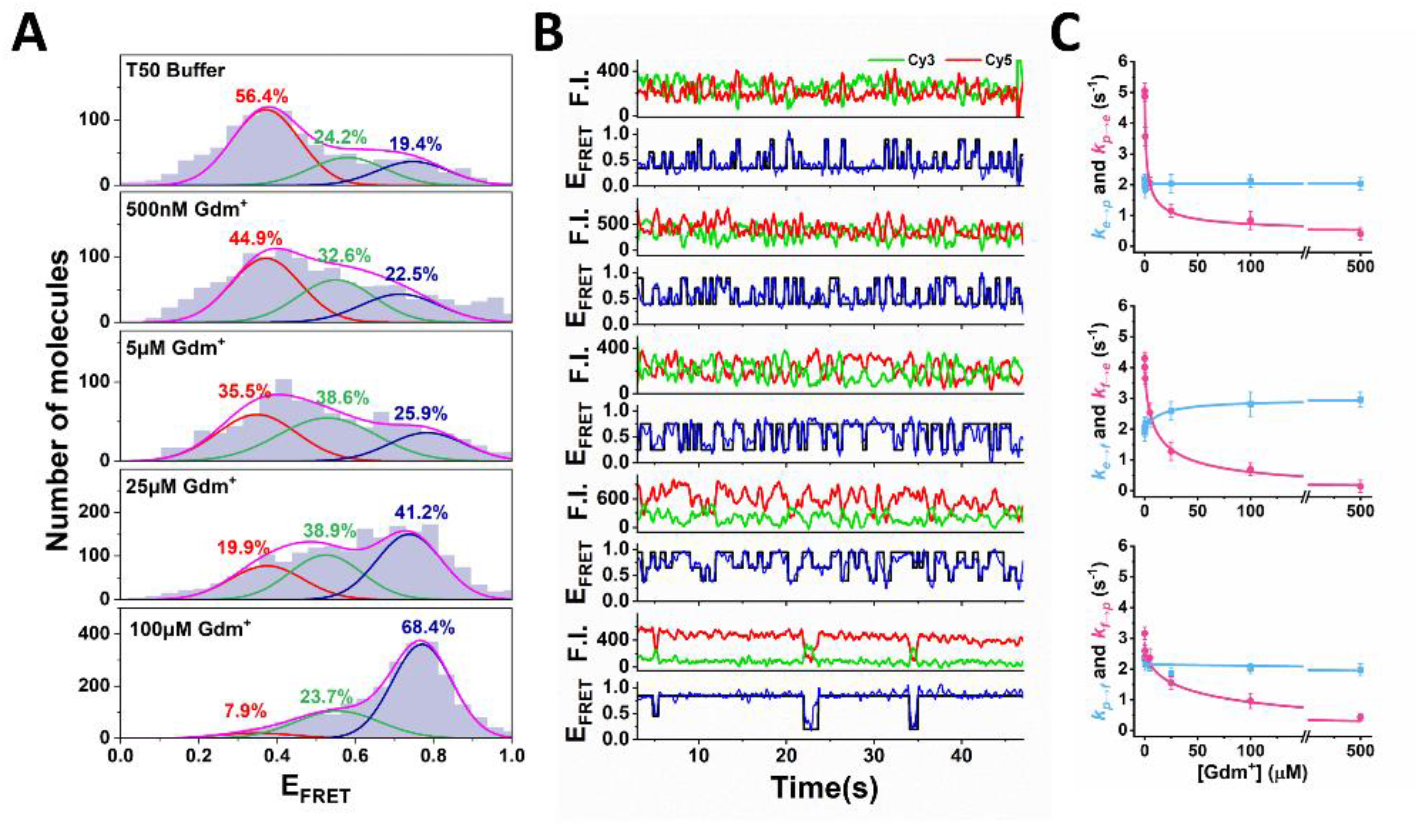
Gdm^+^-dependent folding dynamics of the guanidine-III riboswitch by smFRET. **(A)** FRET population histograms at various Gdm^+^ concentrations. The fraction of three-state from Gaussian fitting is marked in the figure, red for extended state, green for prefolded state and blue for folded state, respectively. **(B)** Representative smFRET traces for guanidine-III riboswitch in different Gdm^+^ concentrations ranging from 0 to 100 µM. The idealized FRET traces generated by the hidden Markov model (black) were overlaid on the experimental traces (blue). **(C)** Transition rate constants for guanidine-III Riboswitch in different Gdm^+^ concentrations.

It is worth noting that no significant change was observed in the forward rates in E-P and P-F transitions with varying Gdm^+^ concentrations, which suggests that conformational selection mechanism takes place in these two transitions(40-42). In contrast, *k*_e→f_ slightly rose from 1.8 s^-1^ to 3.16 s^-1^ at 500 µM Gdm^+^, implying that a proportion of molecules undergoes extended-folded state transition via induced fit(43-45).

### Molecular Dynamics simulations reveal folding intermediate of Guanidine-III riboswitch

To further validate our smFRET experimental findings, we conducted molecular dynamics (MD) simulations using an all-atom structure-based model(46). Our simulations successfully captured the folding pathway of the guanidine-III riboswitch coupled with ligand binding, as demonstrated in Figure 4A. Notably, the two-dimensional free energy landscape analysis (Figure 4B) identified two distinct intermediate states, designated I_1_ and I_2_. Structural analysis revealed that these intermediates were characterized by partial formation of the triplex interactions (Figure 4B, top panel), corresponding to the initial stage of triplex structure assembly(47). Contact number calculations highlighted stark differences between I_1_ and I_2_: the A5-A19 interaction increased from 0.32 ± 0.12 in I_1_ to 4.69 ± 2.31 in I_2_, while A5-G34 contacts surged from 2.06 ± 0.24 to 10.59 ± 7.47. This sharp rise in tertiary interactions indicates that I_1_ represents an extended conformation with partial triplex formation (involving G9, G10, U11 and P2 stem), whereas I_2_ adopts a prefolded state where the ligand-binding pocket is fully preorganized, poised for guanidine recognition.

**Figure 4.**
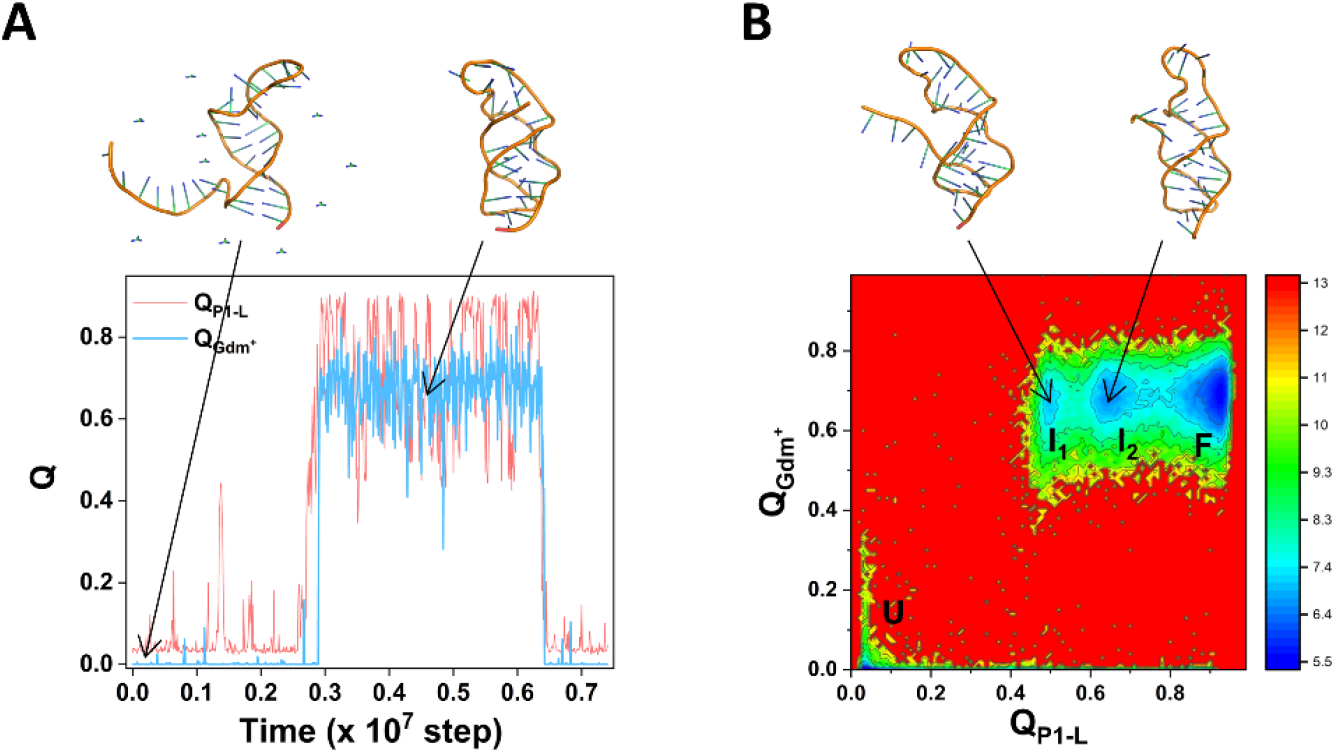
**(A)** A representative MD trajectory illustrating the folding dynamics of guanidine-III riboswitch coupled with guanidine binding. Representative conformations of the unfolded state (left) and folded state (right) shown on the top. **(B)** Two-dimensional free energy landscape projected onto the reaction coordinates of Q^P1-L^ (contact between P1 and L) and Q^Gdm+^ (contact between guanidine-RNA interaction). Representative conformations of the two intermediate states shown on the top.

The simulated intermediates I_1_ and I_2_ align with the E-state (E_FRET_ ≈ 0.35) and P-state (E_FRET_ ≈ 0.6) observed experimentally (Figure 3), validating the hierarchical folding pathway. Importantly, the transition from I_1_ to I_2_ coincides with a ∼15 Å reduction in the A5-G34 distance (Figure 4E), mirroring the FRET efficiency shift from E-to P-state (E_FRET_ ≈ 0.35 → 0.6).

The simulations suggest a triplex-zippering mechanism that aligns with the experimentally observed conformational selection during the E-P transition. In this model, the initial formation of a triplex at the P2 stem (I_1_ state) helps create a partially organized pocket for ligand binding, followed by stepwise helical stacking (I_2_ state) that completes the pseudoknot architecture. Critically, this sequential assembly ensures that only the prefolded intermediates (I_2_) with a structured pocket are capable for Gdm^+^ binding, which is a key feature of conformational selection—where ligands preferentially stabilize already existing structures. The simulated contact dynamics (e.g., A5-G34 interaction surge from 2.06 to 10.59 during I_1_→I_2_) correlated with the Mg^2+^-dependent acceleration of *k*_e→ p_ and *k*_p→ e_ observed experimentally, where Mg^2+^ stabilizes the P-state (I_2_ analog) to favor ligand-competent conformations. This mechanistic synergy explains why the riboswitch predominantly employs conformational selection for E-P transitions: triplex zippering pre-organizes the ligand-binding pocket to enrich the prefolded population.

### Cooperativity between Mg^2+^ and Gdm^+^ on Guanidine-III Riboswitch Folding

Next, to investigate whether the presence of Mg^2+^ enhances the stabilizing effect of Gdm^+^ on the riboswitch structure, we performed Gdm^+^ titration experiments under the conditions of Mg^2+^ concentrations at 10, 50, and 200 µM. The results showed that the riboswitch exhibited a higher and more stable proportion of the F-state at the same Gdm^+^ concentration in the presence of Mg^2+^ (Supplementary Figure S5A, B, C). Notably, when 200 µM Mg^2+^ and 10 µM Gdm^+^ were present simultaneously, the proportion of the folded state reached 90.3% (Supplementary Figure S5C). Transition rate analysis further dissected the interplay between Mg^2+^ and Gdm^+^.

In the presence of Mg^2+^, the forward rates of E-P (*k*_e→ p_) and E-F (*k*_e→ f_) transitions remained largely unperturbed across Gdm^+^ concentrations (Figure 5A–B). Instead, Mg^2+^ reduced the Gdm^+^ concentration required to saturate the folded population, accelerating the riboswitch’s progression from the extended (E) to prefolded (P) and folded (F) states. This suggests that Mg^2+^ pre-organizes the RNA into a conformation primed for ligand binding, lowering the energy barrier for productive folding.

**Figure 5.**
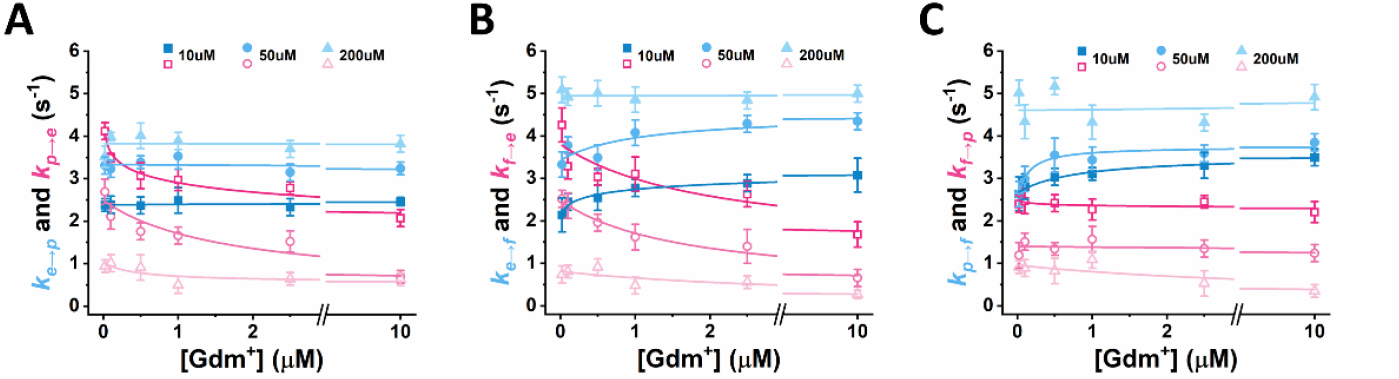
Transition rate constants for guanidine-III riboswitch at different Gdm^+^ concentrations in the presence of 10 µM, 50 µM and 200 µM Mg^2+^. **(A)** extended-prefolded state transitions, **(B)** extended-folded state transitions, and **(C)** prefolded-folded state transitions.

For the P-F transition, distinct Mg^2+^-dependent mechanisms emerged. At low Mg^2+^ (10–50 μM), the reverse rate (*k*_f→ p_) remained constant (∼0.4 s^-1^), while the forward rate (*k*_p→ f_) increased linearly with Gdm^+^ concentration (Figure 5C). This ligand-concentration-dependent acceleration of *k*_p→ f_ —coupled with stable *k*_f→ p_ —aligns with an induced fit mechanism, where Gdm^+^ binding to the partially folded P-state actively reshapes the tertiary architecture to stabilize the F-state. At higher Mg^2+^ (200 μM), however, *k*_p→ f_ plateaued at saturating Gdm^+^ (≥100 µM), indicating Mg^2+^-mediated pre-organization of the pseudoknot core into a ligand-receptive conformation. Under these conditions, the riboswitch transitioned to a conformational selection paradigm, where prefolded intermediates selectively bind Gdm^+^ to lock the final fold.

Our results demonstrate that the presence of Mg^2+^ significantly enhances the response of guanidine-III riboswitch to its ligand, as evidenced by a 5-to 10-fold increase in the slope of the ligand concentration-response curve when plotted on a logarithmic scale (Supplementary Figure S4 and S5). This suggests that Mg^2+^ plays a crucial role in stabilizing ligand-riboswitch interactions, either by directly facilitating ligand binding or by promoting riboswitch conformational changes that enhance ligand sensitivity. The observed increase in slope implies a higher Hill coefficient (Supplementary Figure S5D, E, F), indicating stronger positive cooperativity in ligand binding. Such Mg^2+^-mediated effects likely originate from its capacity to mitigate electrostatic repulsion within the RNA backbone, thereby promoting a more favorable binding environment(48). These results highlight Mg^2+^ as a crucial modulator of riboswitch function, emphasizing the interdependence of divalent cation coordination, ligand sensitivity, and structural switching in metabolite-dependent gene regulation.

### Ligand Binding Specificity of Guanidine-III Riboswitch

To assess the ligand-binding specificity of the guanidine-III riboswitch, we evaluated its conformational response to four other structural analogs of guanidine: aminoguanidine (AG), methylguanidine (MG), metformin (MET), and urea. Under Mg^2+^-free conditions, Gdm^+^ induced a robust 65% increase in the folded fraction (84.4% occupancy at 1 mM Gdm^+^), whereas analogs elicited only modest stabilization (44.4% for AG, 41.3% for MG, and 31.3% for MET at 10 mM; Figure 6A), underscoring the riboswitch’s precision in discriminating Gdm^+^ from analogs lacking its planar geometry or hydrogen-bonding capacity(49).

**Figure 6.**
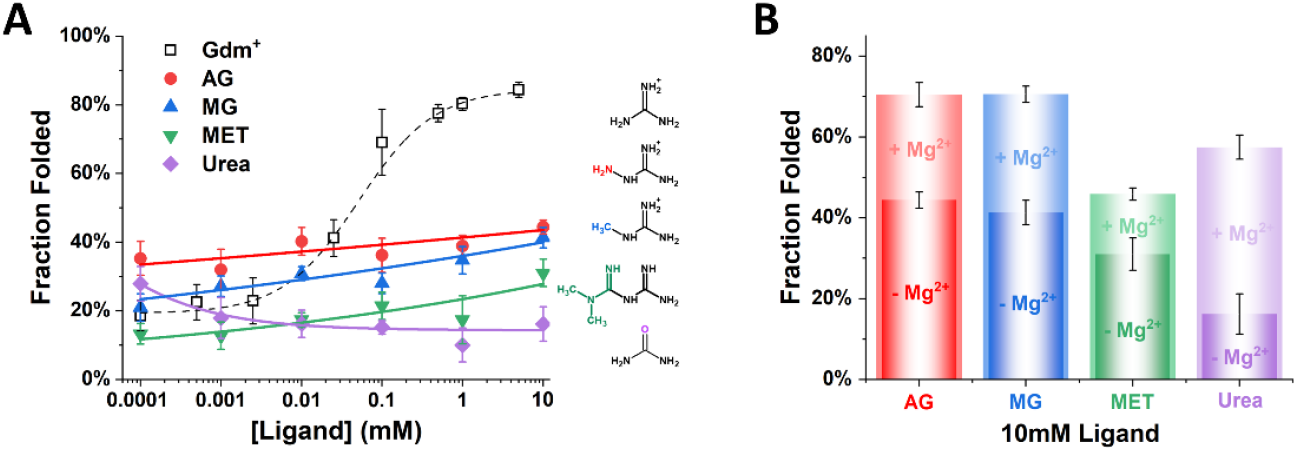
Ligand binding specificity of guanidine-III riboswitch. **(A)** The folded fractions extracted from smFRET population histograms. The data of Gdm^+^ in dashed line was adapted from Supplementary Figure S4B for a comparison. The chemical structures of the four analogs were drawn on the right with color highlighted. **(B)** The folded fractions for each analog at 10 mM concentration in the absence or presence of 1 mM Mg^2+^.

Interestingly, urea exhibited a biphasic effect on riboswitch folding. At concentrations below 1 µM, urea induced a folded fraction of approximately 30%; however, the proportion of the folded state decreased with increasing concentration, reaching 17.3% at 10 mM urea, similar to the observations from in-line probing experiments.(7) This suggests that while urea can transiently stabilize certain riboswitch conformations at low concentrations, it may destabilize the folded state at higher concentrations due to its denaturing effects.

To further explore the interplay between Mg^2+^ and ligand binding, we conducted experiments with 10 mM of each ligand analog in the presence of 1 mM Mg^2+^. The addition of Mg^2+^ increased the folded fraction for all four analogs (Figure 6B), reinforcing our earlier findings that Mg^2+^ facilitates the folding of the guanidine-III riboswitch. These findings highlight the guanidine-III riboswitch’s high specificity for guanidine, while also demonstrating that Mg^2+^ ions can modulate its folding dynamics in the presence of various ligands.

Riboswitches are highly specific RNA elements that regulate gene expression by binding to small metabolites. This specificity is crucial for their regulatory function, as it ensures precise cellular responses to fluctuating metabolite concentrations. The selective binding of riboswitches is fine-tuned by variations in peripheral structural elements that enhance ligand recognition. Similarly, our study demonstrates that the guanidine-III riboswitch exhibits high specificity for guanidine over structurally related analogs. This high specificity likely arises from the precise architecture of its ligand-binding pocket, which is optimized to accommodate the unique structural features of guanidine.

## SUMMARY and CONCLUSION

In this study, we systematically dissected the structural dynamics of the guanidine-III riboswitch using smFRET and MD simulations. Our findings reveal that the riboswitch exhibits three distinct conformational states (extended, prefolded, and folded state) and transitions between these states are regulated by Mg^2+^ and guanidine binding.

Mg^2+^ plays a crucial role in pre-organizing the RNA structure, stabilizing intermediate conformations, and promoting efficient ligand recognition. Guanidine binding primarily suppresses reverse transitions, effectively locking the riboswitch into its folded, active conformation. The interplay between Mg^2+^ and guanidine demonstrates a cooperative mechanism in which Mg^2+^ enhances ligand-induced folding through a combination of conformational selection and induced fit.

These insights contribute to our broader understanding of riboswitch-mediated gene regulation, highlighting how small molecules and ions synergistically modulate RNA structure and function. Given the fundamental role of riboswitches in bacterial physiology, our study provides a framework for potential therapeutic interventions targeting metabolite-sensing RNA elements. Future studies leveraging time-resolved single-molecule approaches and *in vivo* validations will further clarify the mechanistic intricacies of riboswitch dynamics and their regulatory impact on bacterial adaptation.

These findings reinforce a paradigm in riboswitch biology: ligand specificity is achieved not merely through static structural complementarity but via dynamic coupling between ion-mediated folding and ligand-induced kinetic trapping. Such mechanisms ensure that riboswitches respond exclusively to physiologically relevant metabolites, even in environments crowded with structurally similar molecules.

Collectively, these observations underscore the evolutionary adaptation of riboswitches to finely discriminate between metabolites, ensuring accurate regulation of gene expression in response to specific cellular signals.

## SUPPLEMENTARY DATA

Supplementary Figures S1-S6 can be downloaded through website.

## ACKNOWLEDGEMENTS

The authors thank Dr. Qiushi Li at WIUCAS for insightful discussion on the state transitions analysis. This research was funded by the start-up funding from Wenzhou Institute, University of Chinese Academy of Sciences (WIUCASQD2021007, WIUCASQD2022003).

