## Supplemental Figures S1-S6 for "Dynamic Pathway of Guanidine-III Riboswitch Folding Revealed by Single-Molecule FRET: Mg^2+^-Assisted Preorganization and Ligand-Induced Kinetic Trapping"

### Supplementary Figures

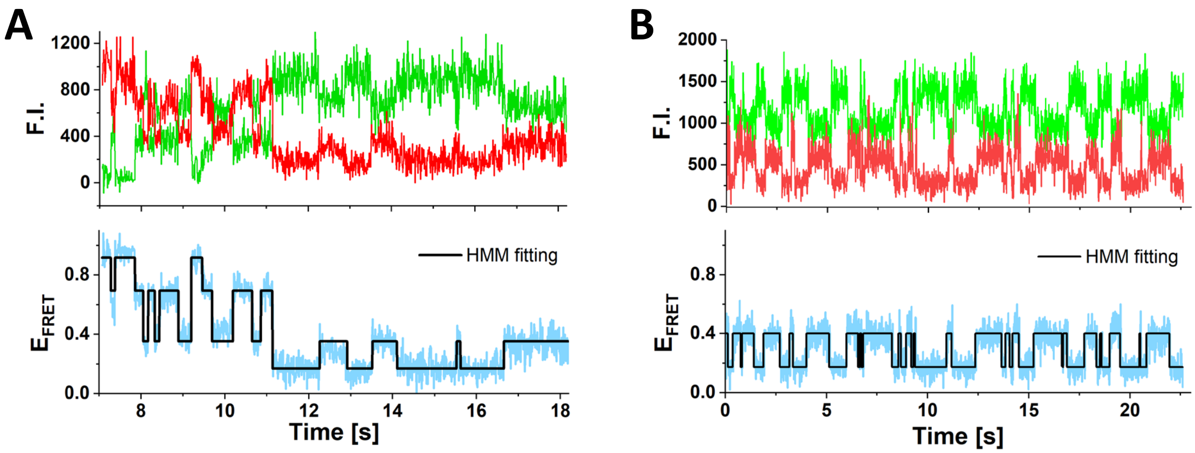


Figure S1. Representative smFRET traces for the unfolded state. **(A)** Trace showing four-state transitions, and **(B)** Transitions between the extended state and the unfolded state. The black lines in the lower panels represent the hidden Markov model fittings.

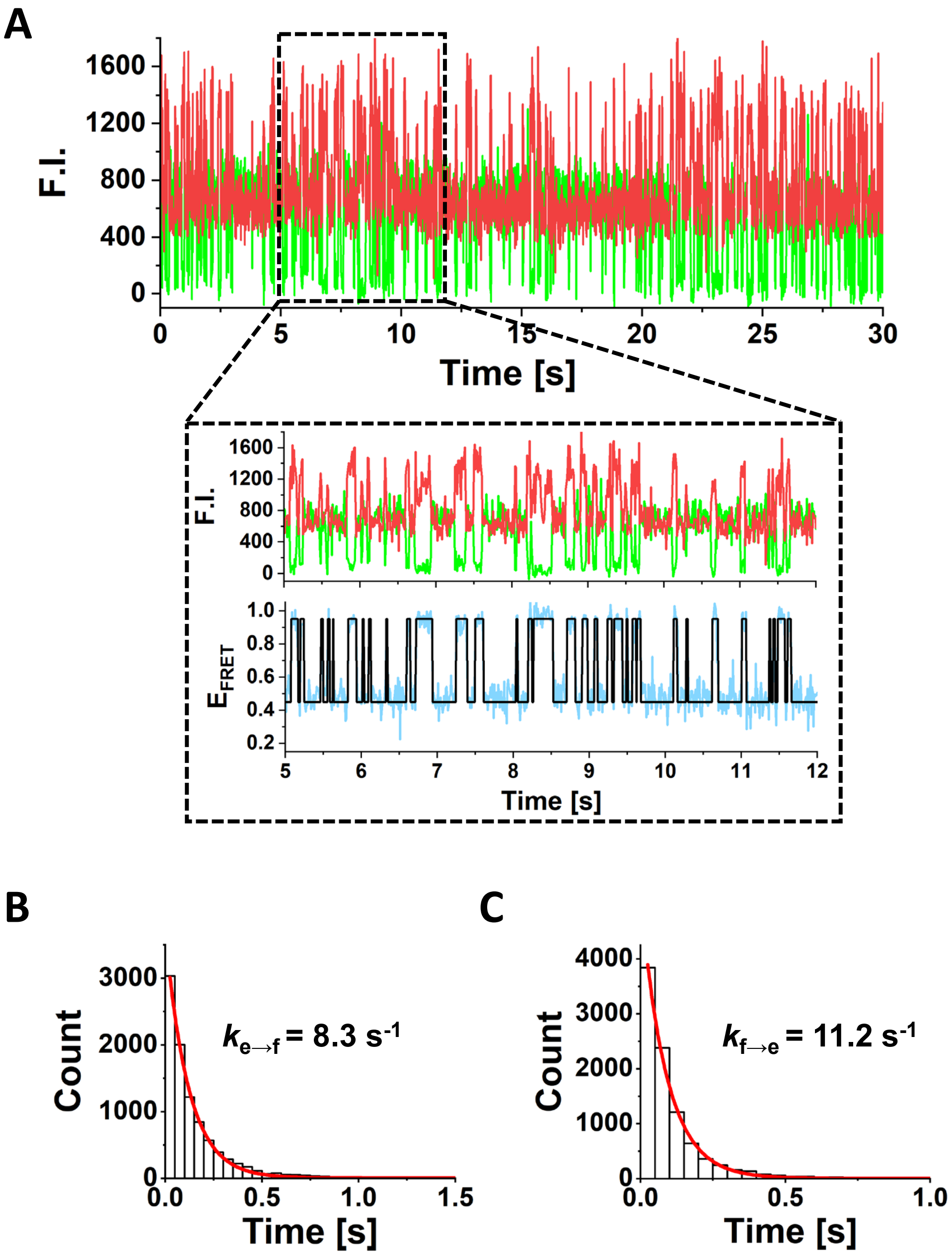


Figure S2. Guanidine-III riboswitch conformational transitions between two states in salt free condition **(A)**. Single-exponential fittings of the dwell time of the extended state and the folded state give the transition rate $k_{e \to f}$ = 8.3 s-1 **(B)** and $k_{f \to e}$ = 11.2 s-1 **(C)**, respectively.

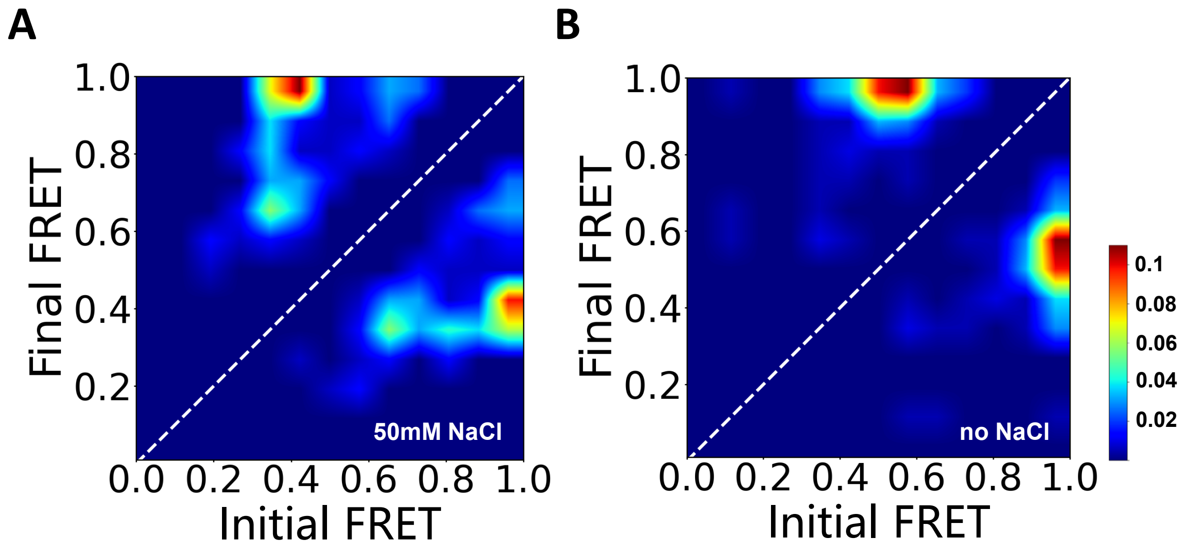


Figure S3. Transition occupancy density plots (TODPs) for guanidine-III riboswitch in the presence **(A)** or absence **(B)** of $Na^{+}$. The plots were generated from smFRET traces of 2-3 independent experiments.

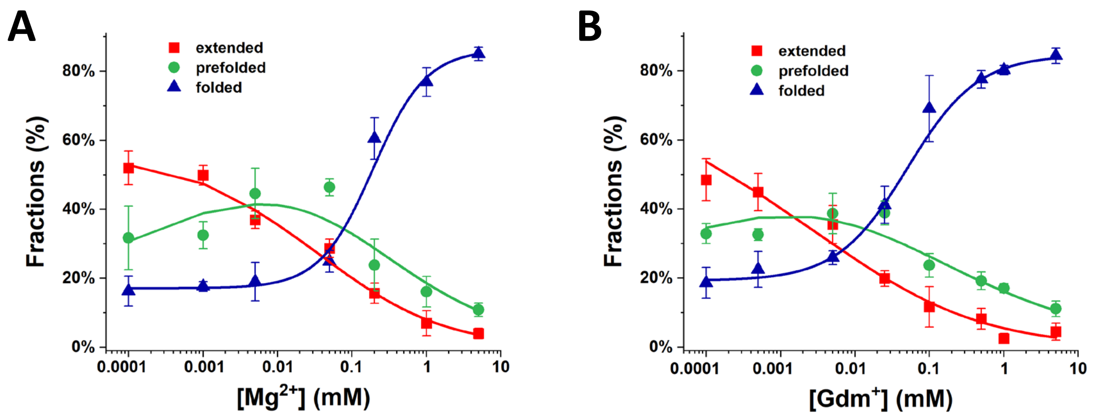


Figure S4. Plots of fraction for each state in the presence of $Mg^{2+}$ **(A)** and $Gdm^{+}$ **(B)**. The factions of folded were fitted with Hill equation to generate the Hill coefficient and $K_D$. The fractions for extended and prefolded state were fitting using a 3-state Hill equation.

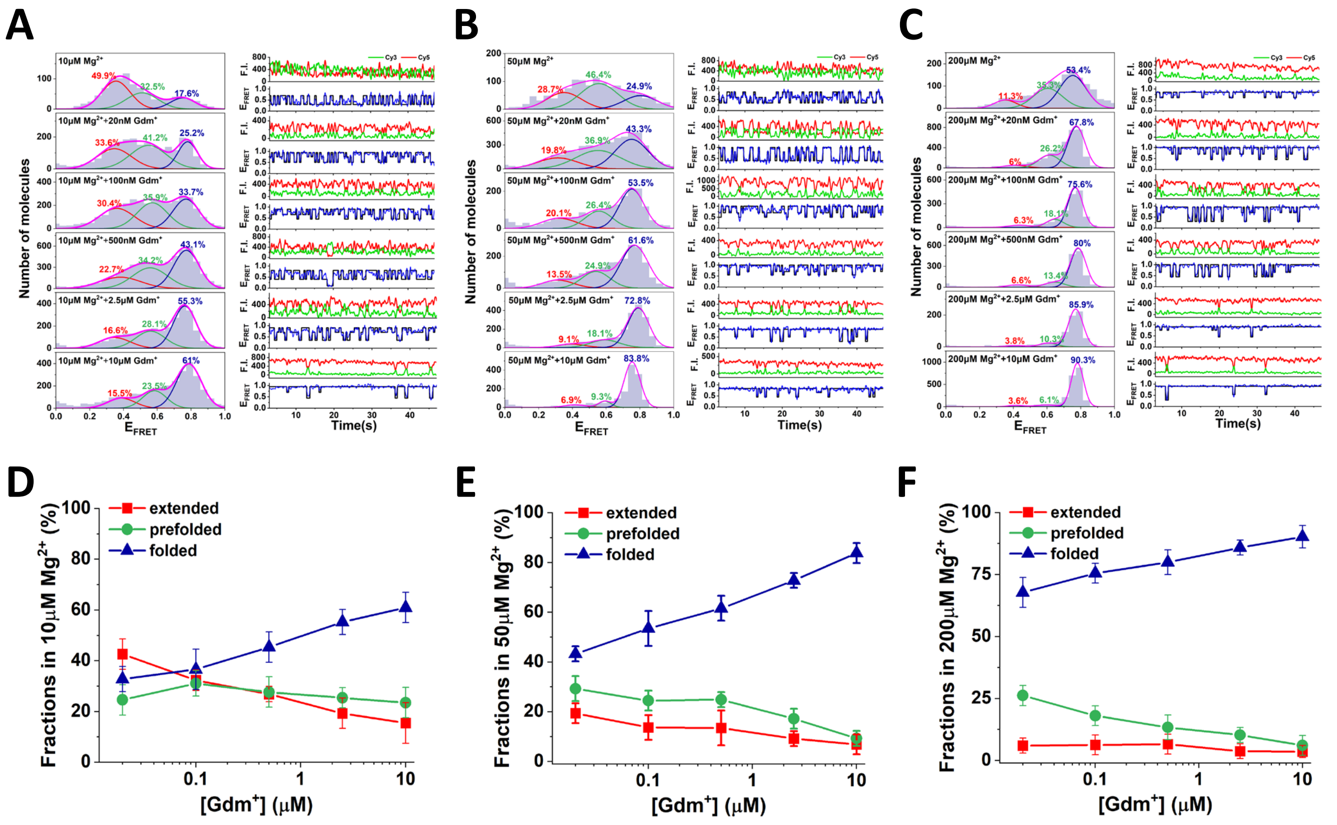


Figure S5. Gdm⁺-dependent folding of the guanidine-III riboswitch in the presence of 10μM $Mg^{2+}$ **(A)**, 50μM $Mg^{2+}$ **(B)** and 200μM $Mg^{2+}$ **(C)**. The histograms of the population of different state were fitted with three-peak Gaussian (left panels). The factions for each states were plotted in **(D)** for 10 μM $Mg^{2+}$, **(E)** for 50 μM $Mg^{2+}$ and **(F)** for 200 μM $Mg^{2+}$, respectively.

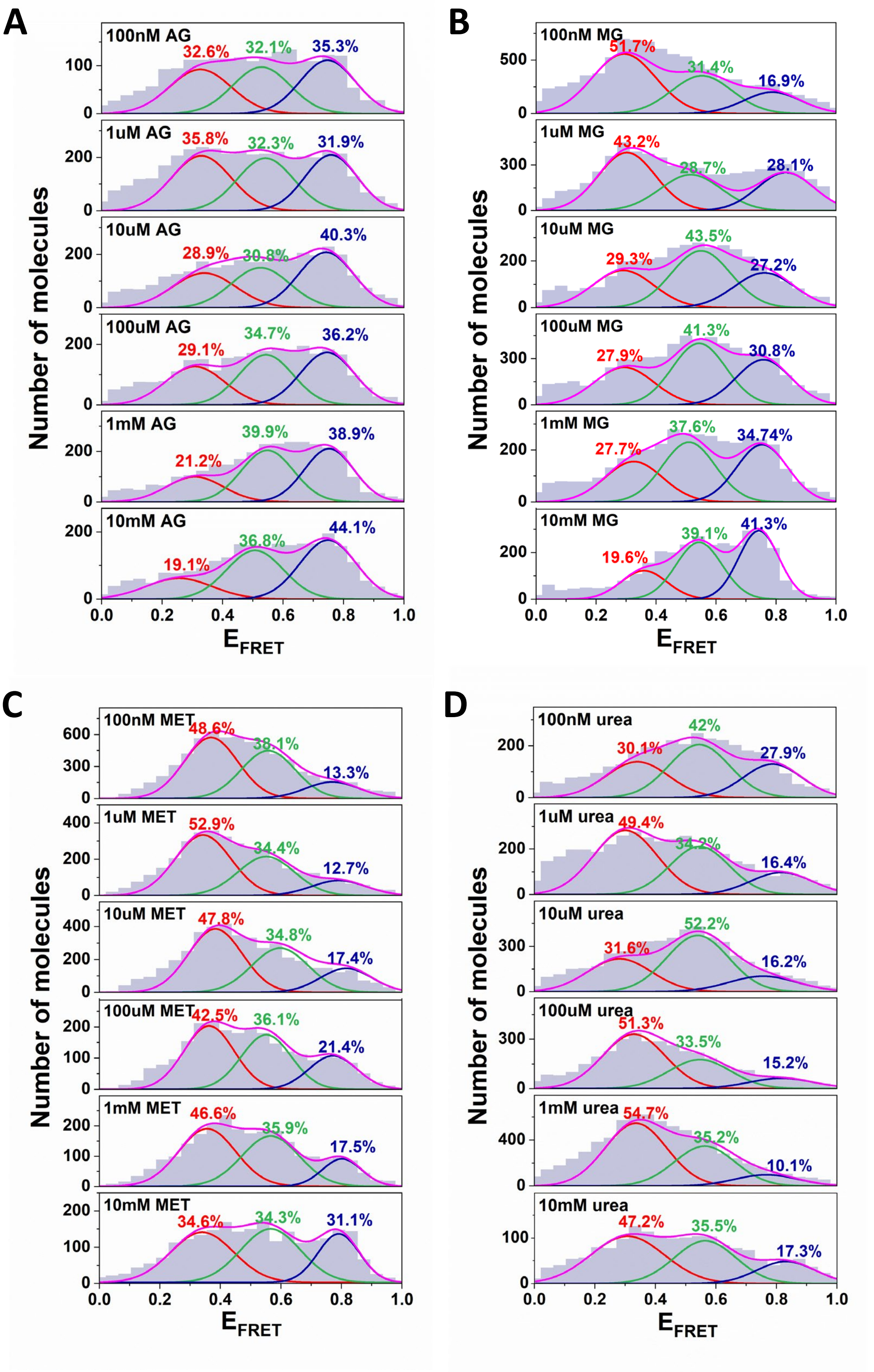


Figure S6. The histograms of the FRET population for **(A)** aminoguanidine (AG), **(B)** methylguanidine (MG), **(C)** metformin (MET), and **(D)** urea. The Gaussian fitting for each state is marked in red for extended state, green for prefolded state and blue for folded state, respectively.
